# RBPSponge: genome-wide identification of lncRNAs that sponge RBPs

**DOI:** 10.1101/566828

**Authors:** Saber HafezQorani, Aissa Houdjedj, Mehmet Arici, Abdesselam Said, Hilal Kazan

## Abstract

**Summary:** Long noncoding RNAs (lncRNAs) can act as molecular sponges or decoys for an RNA-binding protein (RBP) through their RBP binding sites, thereby modulating the expression of all target genes of the corresponding RBP of interest. Here, we present a web tool named RBPSponge to explore lncRNAs based on their potential to act as a sponge for an RBP of interest. RBPSponge identifies the occurrences of RBP binding sites and CLIP peaks on lncRNAs, and enables users to run statistical analyses to investigate the regulatory network between lncRNAs, RBPs and targets of RBPs.

**Availability:** The web server is available at https://www.RBPSponge.com

## 1 Introduction

Eukaryotic genomes encode for thousands of long non-coding RNAs (lncRNAs); however, their functions are largely unknown (Kopp and Mendell, 2018). Recent studies revealed that some lncRNAs can function as microRNA (miRNA) sponges (Mitello *et al*., 2017). These lncRNAs contain several miRNA binding sites and the resulting competition restricts miRNA’s availability to bind to its own targets. In turn, miRNA’s activity is suppressed. A similar relationship has been recently discovered between lncRNAs and RBPs. For instance, a highly conserved cytoplasmic lncRNA, NORAD, contains several functional binding sites for the two mammalian Pumilio homologs (i.e., PUM1 and PUM2) (Tichon *et al*., 2016; Lee *et al*., 2016). NORAD sequesters PUM1/PUM2 molecules and modulates the expression of their target genes. Similarly, Kim et al has been shown that the lncRNA OIP5-AS1 sponges ELAVL1 in HeLa cells (Kim *et al*., 2016). Lastly, Chiu et al performed extensive computational analyses on TCGA datasets where several candidate sponge lncRNAs are predicted for RBPs (Chiu *et al*., 2018).

A number of tools are available to identify lncRNAs that can act as miRNA sponges (Furio-Tari *et al*., 2016). There are also tools to map RBP binding sites on lncRNAs (e.g. Wu *et al*., 2018), however they do not evaluate the enrichment and distribution of binding sites or the regulatory relationship between the lncRNA and RBP target background genes. As such, tools that explore lncRNAs that could function as RBP sponges are still lacking. Here, we introduce a web tool named RBPSponge that explores lncRNAs based on their potential to act as a sponge for an RBP of interest. RBPSponge leverages several types of data such as RBP binding preferences, CLIP datasets and gene expression data. In addition to identifying the occurrences of RBP binding sites on lncRNAs, RBPSponge runs several types of analyses to evaluate the sponge potential of lncRNAs.

## 2 Data and Methods

Human RBPs with known existing binding preferences are compiled by merging position weight matrices (PWMs) from three compendiums: RNAcompete (Ray *et al*., 2013), RBNS (Dominguez *et al*., 2017) and RBPmap (Paz *et al*., 2014). These RBPs are further filtered by selecting those with CLIP data compiled from ENCODE eCLIP datasets (van Nostrand *et al*., 2016) and CLIPdb database (Yang *et al*., 2015) (See Supplementary Table 1 for the complete list of CLIP datasets). 40 RBPs are retained after these steps.

### 2.1 Identifying potential lncRNAs that sponge RBPs

The top 3 scoring k-mers were determined for each PWM to represent the set of binding motifs for the RBP. For RBPs with multiple PWMs, the union of top 3 scoring k-mers is used as the set of binding motifs. lncRNA sequences (exon regions only) that are downloaded from GENCODE database (v25lift37) are scanned for k-mer occurrences. To evaluate the significance of RBP motif occurrences in lncRNA sequences two metrics are calculated: log-odds (LOD) score and dispersity score (Furio-Tari *et al*., 2016). LOD score evaluates the enrichment of RBP binding sites on a lncRNA. To obtain the LOD score for a lncRNA/RBP pair, we calculate the number of motif occurrences across each sliding window (of size *w*) of the lncRNA sequence. Assuming that *n_t,w,i_* corresponds to the number of occurrences of binding motifs within the window that starts at position *i* of lncRNA *t*, 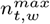 is defined as *max*(*n*_*t,w*,1_, *n*_*t,w*,2_,…,*n_t,w,l_*) where *l* is the starting position of the last sliding window within the lncRNA. The sliding window approach enables the identification of the window with maximum number of motif occurrences. 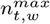 is then normalized by the average maximum number of occurrences of the same set of binding motifs across all lncRNAs :

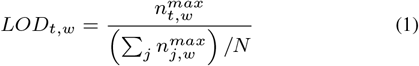

where N is the number of all lncRNAs. LOD score is calculated with varying window sizes (from 50nts to 1000nts in steps of 50nts) and the window size with the largest LOD score is reported. Avoiding the use of a fixed window size enables a robust motif enrichment analysis for lncRNAs of varying length. LncRNAs with high LOD scores contain an enrichment of binding sites for the RBP of interest compared to other lncRNAs (Figure S2).

The second metric which is named dispersity score evaluates the clustering of motif occurrences within the sequence. To calculate the dispersity score of an lncRNA/RBP pair, we build a vector of normalized 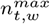 values across all window sizes :

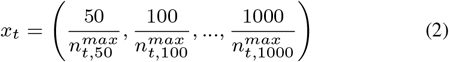

Then, the dispersity score for lncRNA *t* is calculated as the standard deviation of vector **x_t_**. Smaller dispersity scores correspond to a more even distribution of motif occurrences across the sequence. Equal distribution of motifs are observed for lncRNAs that sequester miRNAs (Memczak *et al*., 2013) and we hypothesize that lncRNAs that sequester RBPs show a similar property. In summary, LOD score focuses on the number of binding sites whereas dispersity score focuses on the distribution of these binding sites (Figure S2).

We define threshold values for the LOD / dispersity scores by finding the scores at the 95% percentile of the distribution of all possible RBP-lncRNA pairs. This resulted in the values 1.6 and 36 for LOD score and dispersity score, respectively.

To incorporate experimental binding data, CLIP peaks that are located within lncRNAs are determined. Lastly, gene expression datasets are compiled to assess the regulatory network between lncRNAs, RBPs and their target genes. The following datasets are used: (i) GTEX; (ii) E-MTAB-2706; (iii) E-MTAB-2770. Utilizing all these resources, the following information is displayed for each RBP-lncRNA pair (Figure 1):

- number of non-overlapping motif occurrences in the entire lncRNA sequence
- number of non-overlapping motif occurrences within the CLIP peaks that are located in the lncRNA
- log-odds enrichment score of motif frequencies
- dispersity score
- number of eCLIP / CLIPdb peaks
- median and maximum expression value across the samples available in GTEX, E-MTAB-2706, E-MTAB-2770 datasets
- consensus score

The consensus score is calculated by counting the number of satisfied constraints listed below:

- LOD score > 1.6
- dispersity score < 36
- existence of at least one motif in eCLIP or CLIPdb peak
- maximum expression > 5 TPM in at least one dataset
- correlation analysis gives a p-value < 0.05 in at least one dataset
- regression analysis gives a p-value < 0.05 in at least one dataset

**Fig. 1.**
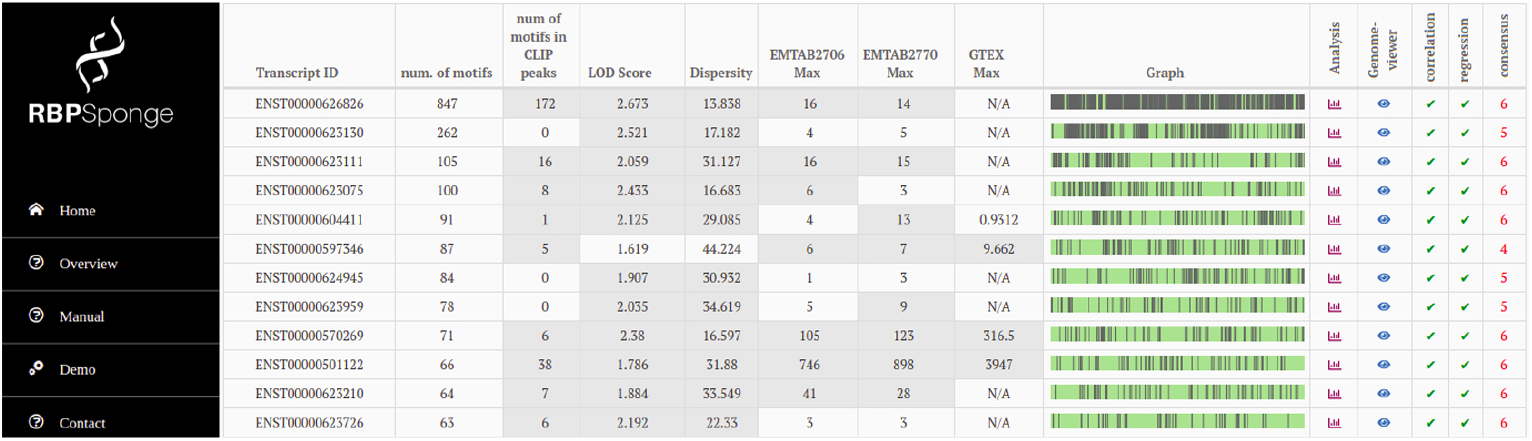
Screenshot from RBPSponge website. Putative sponge lncRNAs for PUM2 are listed in a table where columns display information on motif occurrences and CLIP peaks.

As each positive check increases the consensus score by 1, consensus scores range between 0 and 6. The results are displayed in decreasing order of consensus scores.

## 3 Identifying target and background gene sets for each RBP

Target and background genes sets of RBPs are needed for the statistical analyses implemented by RBPSponge. The user can either upload these gene sets or use our pre-defined gene lists. To define these lists of genes for each RBP, gene annotation files are downloaded from GRCh37 assembly of Ensembl (Release 87) and the longest 3’UTR isoform is determined for each gene. All 3’UTRs are scanned with the set of k-mers that are determined for each RBP. 3’UTRs are also intersected with CLIP peaks to determine overlaps as the presence of a CLIP peak strongly suggests that the region is bound by the RBP of interest.

Data from shRNA knockdown assays are downloaded from ENCODE project (i.e. see Supplementary Table 1 for the complete list). For these assays, log fold changes (LFCs) are calculated using DeSeq2 method Love *et al*. (2014). For ELAVL1, knockdown dataset from Mukharjee et al Mukherjee *et al*. (2011) is used as knockdown assay data for ELAVL1 was not available in ENCODE project.

We defined the target set as those genes that satisfy the following conditions: (i) occurrence of at least one CLIP peak; (ii) occurrence of one of the top 3 scoring k-mers within a CLIP peak; (iii) has an LFC value that has an adjusted p-value < 0.05 as calculated by DeSeq2 for knockdown datasets.

Similarly, we defined the background set as those genes that satisfy the following conditions: (i) no CLIP peak occurrence; (ii) no occurrence of top 3 scoring k-mers; (iii) significant LFC value in knockdown datasets. We also ensured that the background genes are length-matched to target genes such that there is no statistically significant difference in distribution of length between target set and background set.

## 4 Analysis

The expression values of the lncRNA and the RBP across the samples of the selected dataset are displayed with bar plots. This is an important sanity check as most lncRNAs have low abundance compared to other molecules in the cell and this might prevent them to act as molecular decoys for RBPs. In addition to this, three types of analyses are performed (scripts are available on github page: https://github.com/rbpsponge/RBPsponge).

In the first analysis, we investigate how the expression of the lncRNA co-vary with the expression of target genes of the RBP. To assess the distribution of correlation coefficient values that we obtain with target genes, we repeat the same analysis with a set of background genes as a control. If the lncRNA acts as a decoy for the RBP of interest, RBP activity is reduced and the expression levels of its target genes are affected. In summary, we calculate the Spearman correlation coefficient between the expression values of the lncRNA and the expression values of each gene in the target/background set. We compare the distribution of correlation values of target and background genes with Wilcoxon rank-sum test and also display them with a box plot.

In the second analysis, we assess whether lncRNA expression has added predictive value in addition to RBP expression in determining the expression levels of target genes. If the lncRNA affects the activity of the RBP as a decoy, including lncRNA expression should improve the performance. To this end, we perform a simple linear regression analysis to predict target gene expression where in one case only RBP expression is used and in the other case both RBP and lncRNA expression are used as features. We calculate the Spearman correlation values between actual and predicted expression values of target genes on held-out datasets using 10-fold cross-validation. If the lncRNA of interest acts as a sponge for the RBP, we expect to see an improved predictive performance when lncRNA expression is included as an additional feature. The significance of change is evaluated with Wilcoxon rank-sum test and likelihood-ratio test.

In the third analysis, we look into the expression changes upon the knockdown of the lncRNA of interest, when available. Because lncRNA activity is minimized we expect to see an increased RBP activity and a more pronounced effect (either stabilizing or de-stabilizing) on the expression of target genes. To this end, we compare the distribution of expression changes of target and background genes with a cumulative distribution frequency (CDF) plot. Wilcoxon rank-sum test is used to assess the significance between the two distributions.

## 5 Example run

As input, user selects an RBP of interest and optionally uploads a set of target and background genes for this RBP. As output, a table is displayed where each row corresponds to a lncRNA and columns display the number of motifs, number of CLIP peaks etc. (as described in Section 2.1, Figure 1A). The graphical representation on the right part displays the positions of motif occurrences within that lncRNA as vertical bars. Motifs and CLIP peaks can be also explored within an integrated genome viewer that is displayed as a pop-up window (Supplementary Figure 1).

When the analysis button is clicked, a multi-tab page is displayed where each tab corresponds to a different analysis. For example, when E-MTAB-2706 dataset is chosen for PUM2 (RBP)-NORAD (lncRNA) pair, expression values of PUM2 and NORAD across the tissues are displayed in the first and second tab, respectively (Supplementary Figure 2A-B). In the third tab, the box plot shows that NORAD expression is correlated higher with target genes compared to the background genes (Supplementary Figure 2C). In the fourth tab, we observe that including NORAD in addition to PUM1/PUM2 expression improves the prediction performance significantly (Supplementary Figure 2D). Lastly, the CDF plot in the last tab shows that PUM2 target genes are stabilized more upon the knockdown of NORAD (Supplementary Figure 2E).

## Funding

This work was supported by European Union [FP7 Marie Curie CIG grant 631986 to HK].

