## Supplementary Material for "RBPSponge: genome-wide identification of lncRNAs that sponge RBPs"

### 1 Gene expression datasets

We use three gene expression datasets to investigate the relation between the expression of RBPs, lncRNAs and target/nontarget genes of the RBPs. The details of these datasets are provided below:

- EMTAB2706: This dataset contains gene expression measurements across 675 human cancer cell lines. A full list of the cell lines can be found in [2]. We used the processed data provided by the authors in https://www.ebi.ac.uk/arrayexpress/experiments/E-MTAB-2706/files. In addition to using all cancer cell lines, we allow tissue-specific analysis for the following cancer types that have more than 40 samples: lung, lymphoid, breast, ovary, skin and colorectal.
- EMTAB 2770: This dataset provides the gene expression measurements for 934 human cancer cell lines. We provide tissue-specific analysis for the following types that have more than 40 samples: lung small cell, breast, melanoma, colorectal, lung NSC, pancreas, glioma and ovary.
- GTEx: This dataset encompasses gene expression measurements for 8555 samples across 31 tissue types. GTEx V7 data release is downloaded from https://gtexportal.org. Tissue-specific analysis is provided for all tissue types except Bladder, Bone Marrow, Fallopian Tube and Cervix Uteri as these have less than 40 samples.

All the datasets mentioned above have TPM values.

### 2 Analysis of Chiu et al results

Chiu et al predicts lncRNA-RBP pairs where lncRNA modulates the activity of the RBP [1]. In addition to lncRNAs that act as decoys, this study also predicts lncRNAs that act as co-factors, guides or switches. As such, we went through the top 100 modulator lncRNA-RBP interactions for each cancer type and identified the pairs where lncRNA contains at least one occurrence of RBP binding motif in a CLIP peak and ncRNA expression is at least 5 TPM in one of the gene expression datasets. We display the analyses results for a subset of these pairs in Figure S5-S7.

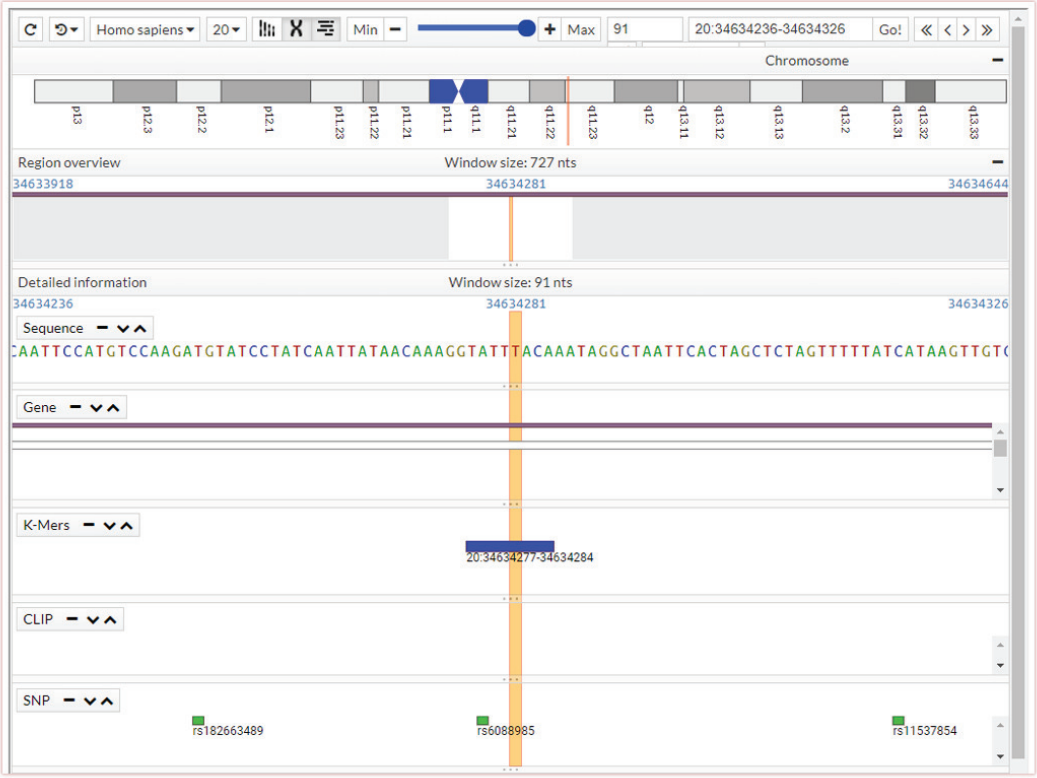


Figure S1: The occurrences of motifs and CLIP peaks within a lncRNA can be explored within an integrated genome viewer [3].

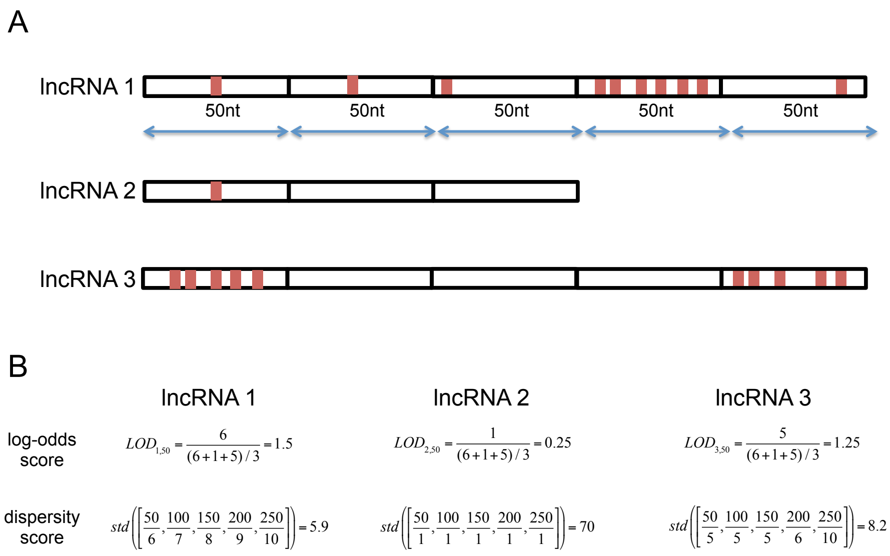


Figure S2: A) A toy example with three lncRNAs to illustrate the log-odds (LOD) and dispersity scores. All lncRNAs are 250nts long and each red bar corresponds to an RBP binding site. LncRNA 1 and lncRNA 3 have 10 binding sites whereas lncRNA 2 has a single binding site. B) LOD and dispersity scores for the three lncRNAs. For LOD score, results for W = 50 are calculated. LncRNA 1 has the highest enrichment for RBP binding sites. Similarly, the binding sites in lncRNA1 are well separated compared to the other two lncRNAs. These are reflected in the scores i.e., lncrna 1 has the highest LOD score and lowest dispersity score.

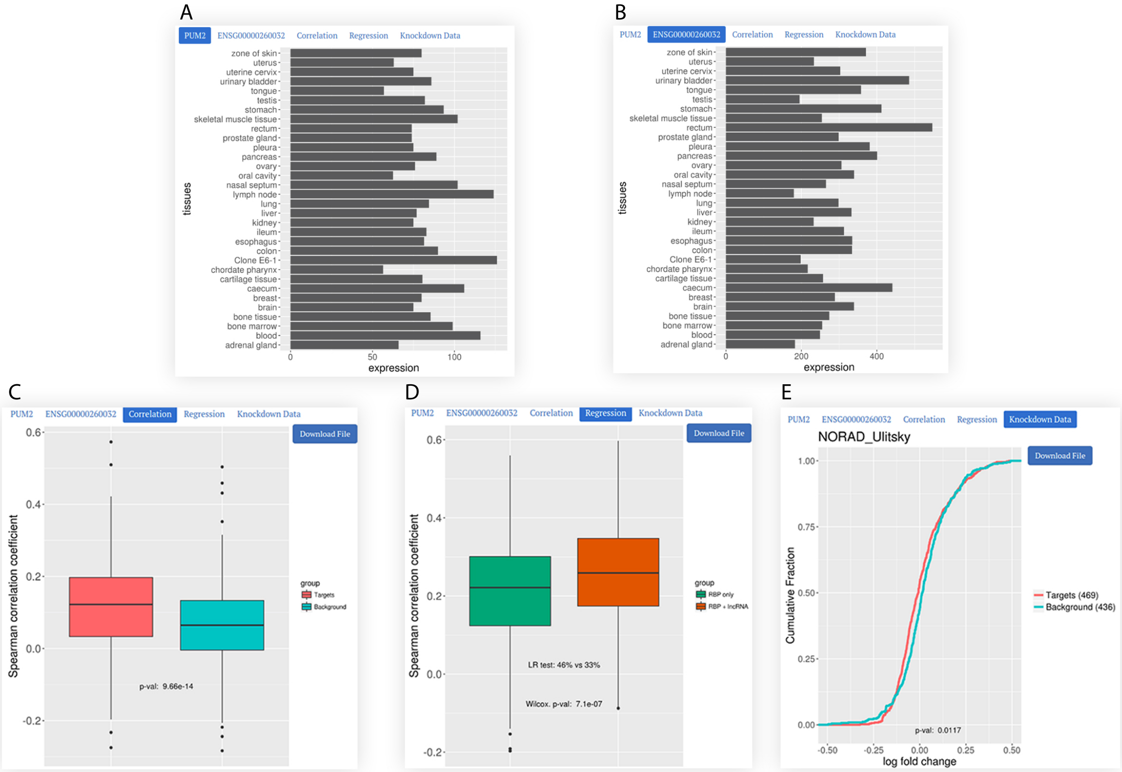


Figure S3: A) Expression values of RBP and NORAD are displayed for the selected dataset (EMTAB-2706). B) Correlation values of NORAD expression with target and background gene expression values are displayed with a box plot. C) Predictive performance of regression models are displayed with a box plot. D) The log fold changes of target and background genes are compared with a CDF plot when NORAD is depleted.

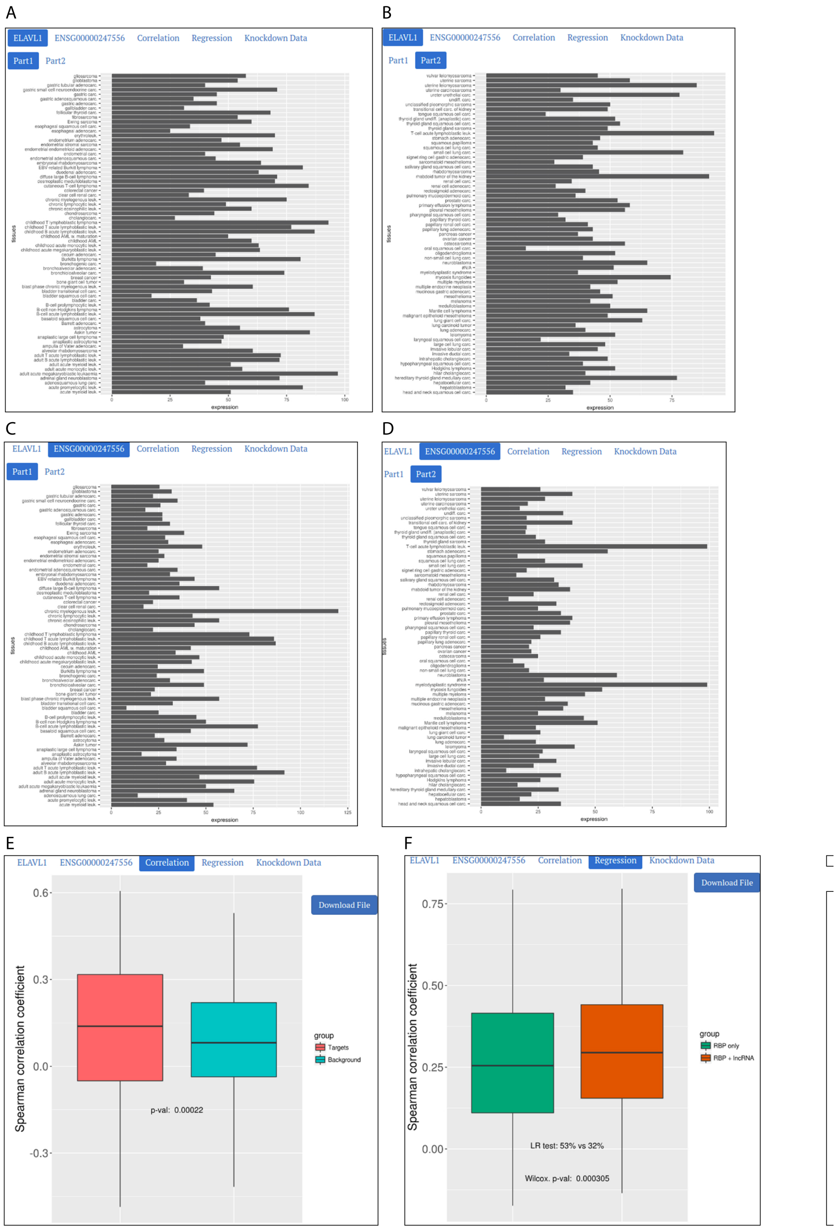


Figure S4: A) Expression values of ELAVL1 and OIP5-AS1 are displayed for the selected dataset (EMTAB-2770). B) Correlation values of OIP5-AS1 expression with target and background gene expression values are displayed with a box plot. C) Predictive performance of regression models are displayed with a box plot.(Knockdown data is not available for this lncRNA.)

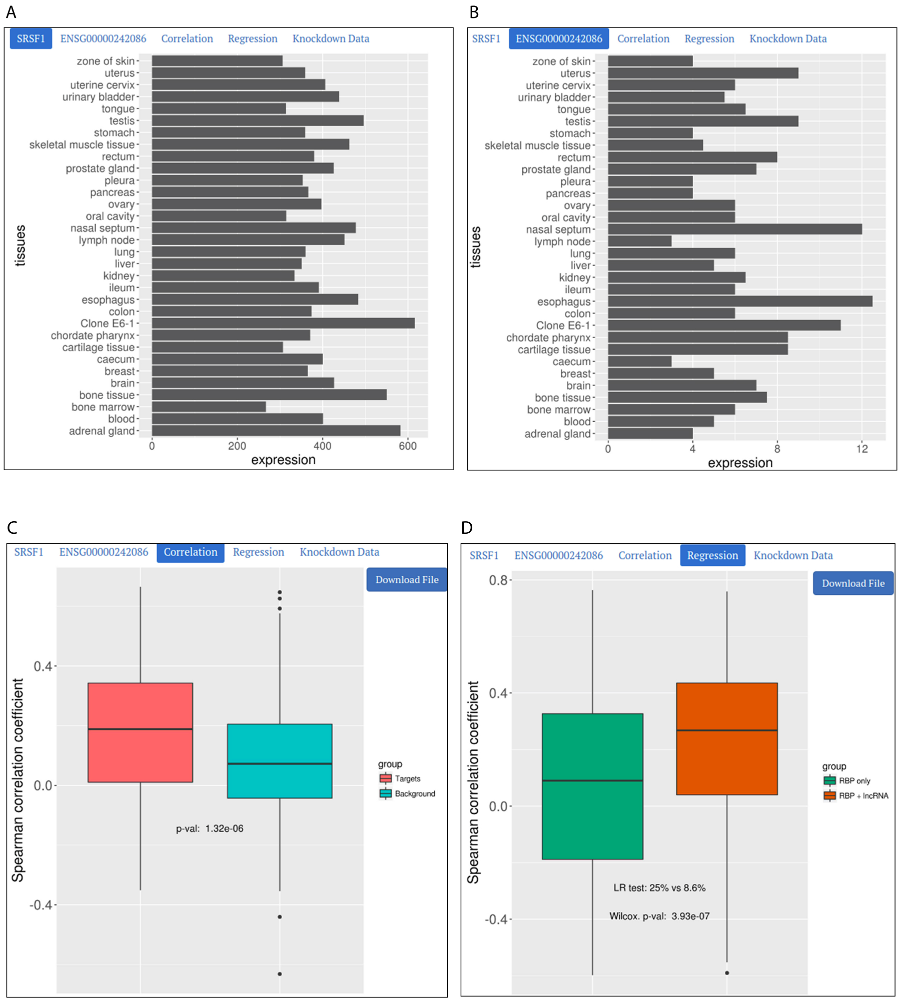


Figure S5: A) Expression values of SRSF1 and LINC00969 are displayed for the selected dataset (EMTAB-2706 skin tissue samples only). B) Correlation values of LINC00969 expression with target and background gene expression values are displayed with a box plot. C) Predictive performance of regression models are displayed with a box plot.(Knockdown data is not available for this lncRNA.)

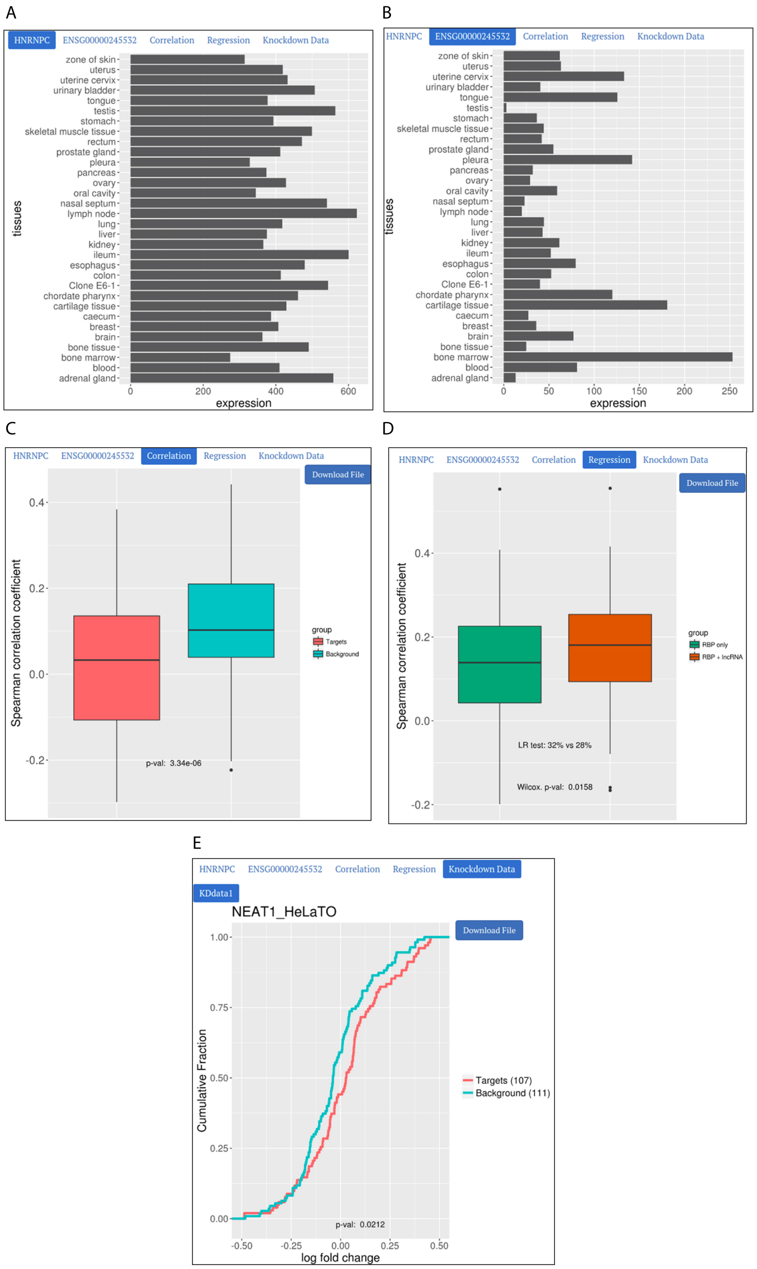


Figure S6: A) Expression values of HNRPC and NEAT1 are displayed for the selected dataset (EMTAB-2706). B) Correlation values of NEAT1 expression with target and background gene expression values are displayed with a box plot. C) Predictive performance of regression models are displayed with a box plot. D)The log fold changes of target and background genes are compared with a CDF plot when NEAT1 is depleted in HeLa cells.

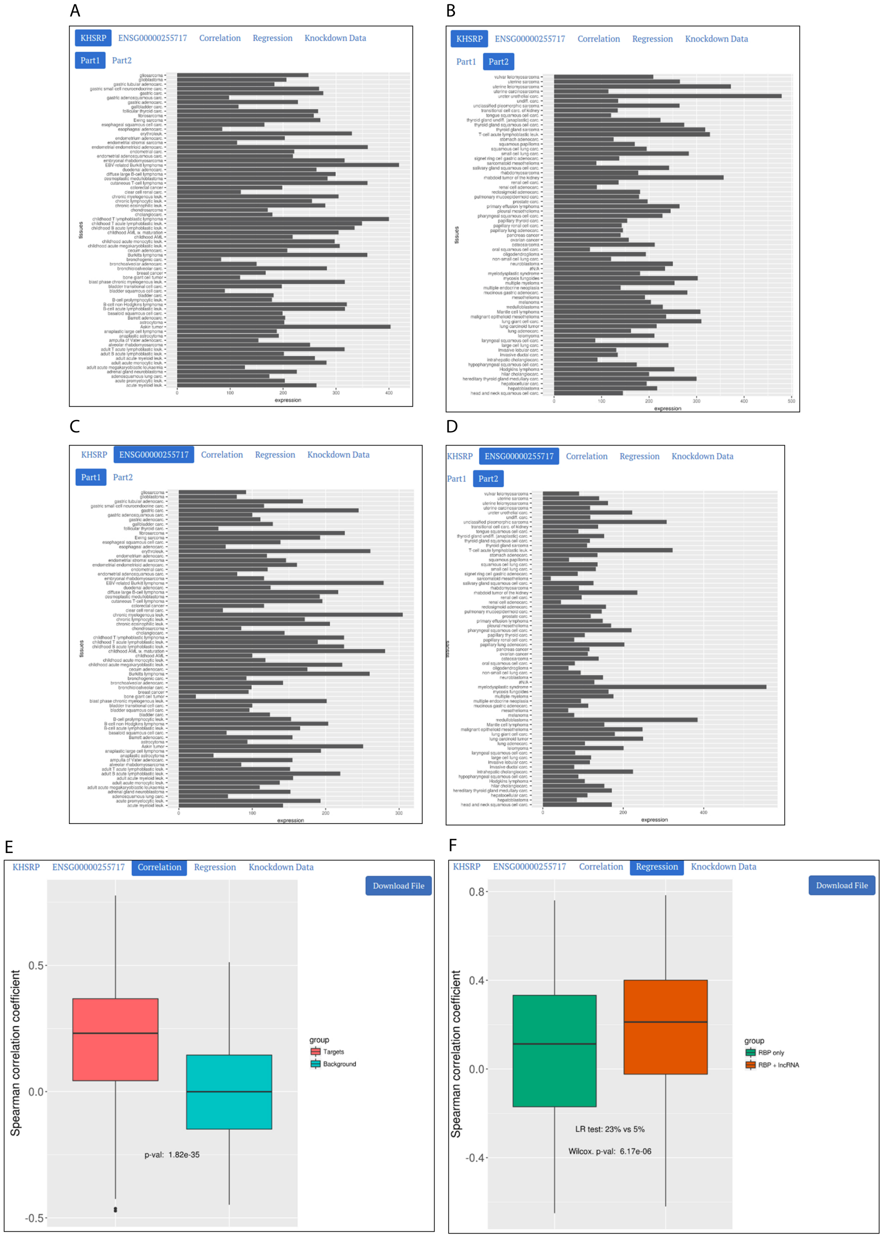


Figure S7: A) Expression values of KHSRP and SNHG1 are displayed for the selected dataset (EMTAB2770 ovarian cancer samples only). B) Correlation values of SNHG1 expression with target and background gene expression values are displayed with a box plot. C) Predictive performance of regression models are displayed with a box plot.(Knockdown data is not available for this lncRNA.)
